# Phosphorylation controls spatial and temporal activities of motor-PRC1 complexes to complete mitosis

**DOI:** 10.1101/2023.03.11.531660

**Authors:** Agata Gluszek-Kustusz, Benjamin Craske, Thibault Legal, Toni McHugh, Julie P.I. Welburn

## Abstract

During mitosis, spindle architecture alters as chromosomes segregate to daughter cells. The microtubule crosslinker Protein Required for Cytokinesis 1 (PRC1) is essential for spindle stability, chromosome segregation and completion of cytokinesis, but how it recruits motors to the central spindle to coordinate the segregation of chromosomes is unknown. Here, we combine structural and cell biology approaches to show that the human CENP-E motor, which is essential for chromosome capture and alignment by microtubules, binds to PRC1 through a conserved hydrophobic motif. This binding mechanism is also used by Kinesin-4 Kif4A:PRC1. Using *in vitro* reconstitution, we demonstrate that CENP-E slides antiparallel PRC1-crosslinked microtubules. We find that the regulation of CENP-E -PRC1 interaction is spatially and temporally coupled with relocalization to overlapping microtubules in anaphase. Finally, we demonstrate that the PRC1:microtubule motor interaction is essential in anaphase to control chromosome partitioning, retain central spindle integrity and ensure cytokinesis. Taken together our findings reveal the molecular basis for the cell cycle regulation of motor-PRC1 complexes to couple chromosome segregation and cytokinesis.

## INTRODUCTION

In cell division, the spindle has a crucial role in ensuring chromosomes are correctly partitioned into daughter cells. Antiparallel microtubules are essential for bipolar spindle stability during mitosis. In anaphase, sister chromatids are pulled to opposite poles. During that stage, re-modelling and elongation of the spindle reduces the risk of DNA damage to lagging chromosomes and aneuploidy, by moving chromosomes away from the cleavage plane. They are physically separated by the central spindle, or midzone, which is a stable structure of antiparallel microtubules that specifies the plane of division.

At the start of mitosis, CENP-E localizes to unattached kinetochores, where it associates with BubR1 and the outer corona (a fibrous expanded structure) of chromosomes (Ciossani et al., 2018; Cooke et al., 1997; Legal et al., 2020; Yen et al., 1991). Kinetochore-bound CENP-E enables kinetochore capture and lateral attachment to microtubules. CENP-E moves chromosomes along the spindle to the metaphase plate before kinetochore biorientation (Fig 1A), reviewed in (Craske et al., 2022). The CENP-E motor relocalizes from kinetochores to the central spindle, at the metaphase to anaphase transition (Kurasawa et al., 2004; Yao et al., 1997). This relocalization is dependent on PRC1 (Protein required for cytokinesis 1), a non-motor microtubule binding protein essential for central spindle assembly. CENP-E has a substantial role in chromosome alignment in early mitosis, but its PRC1-dependent recruitment from kinetochores to the central spindle, during the metaphase to anaphase transition, is less well understood. Depletion or inhibition of CENP-E results in accumulation of mis-attached polar chromosomes, and spindle checkpoint arrest in metaphase, making studies on CENP-E function in anaphase challenging (Chan et al., 1999; Chan et al., 1998; Qian et al., 2010; Schaar et al., 1997). Small molecule inhibition of CENP-E in anaphase and telophase results in spreading and delocalization of PRC1 on the central spindle. This observation led to a proposed role for CENP-E in organizing overlapping microtubules and the central spindle (Liu et al., 2020).

**Figure 1:**
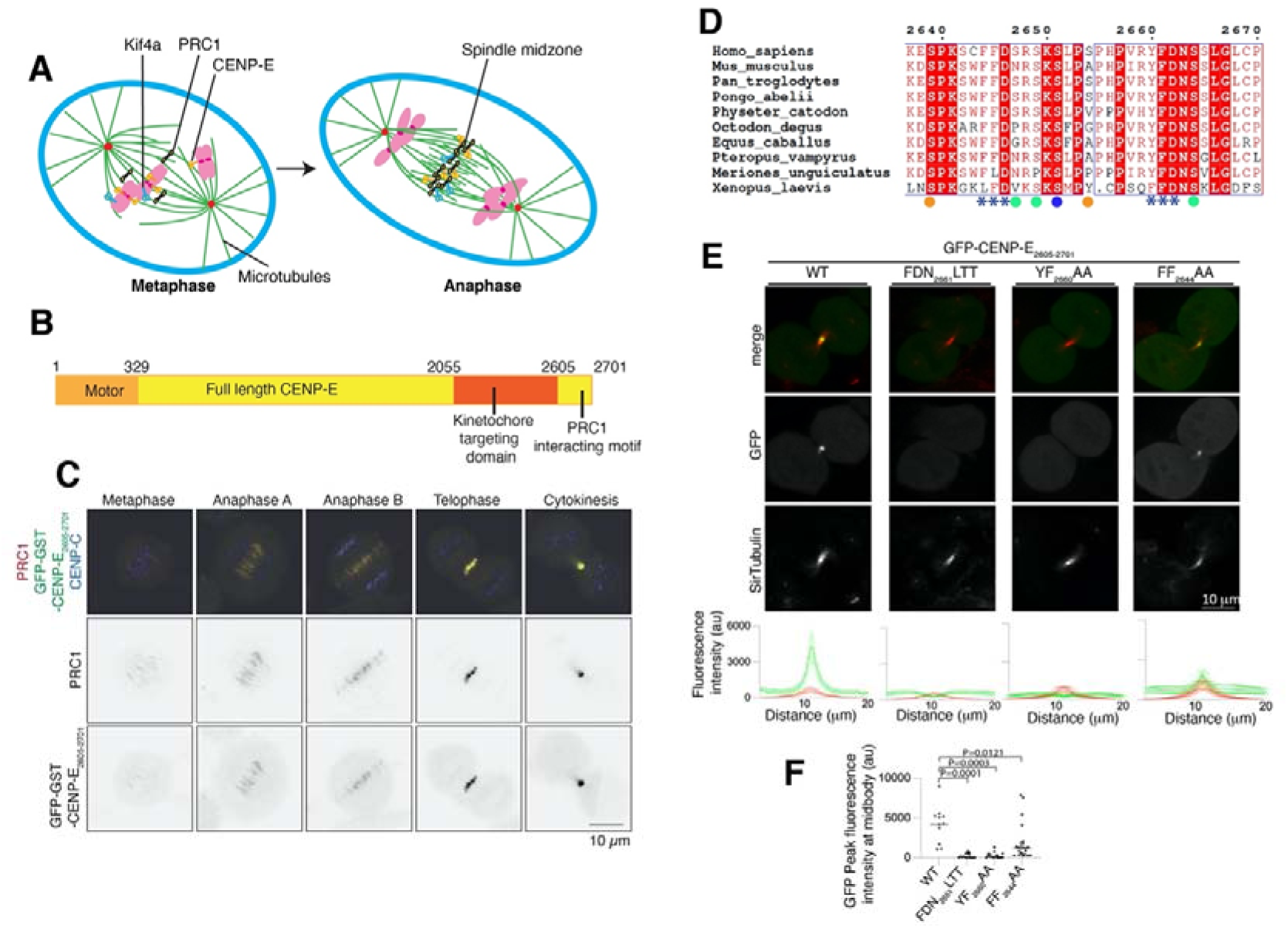
A hydrophobic motif is essential for recruitment of CENP-E to overlapping microtubules in mitosis. (A) Schematic diagram showing the metaphase to anaphase transition, during which kinesin motors Kif4a (blue) and CENP-E (orange) relocalize from chromosomes and kinetochores (pink) to PRC1 (black) on crosslinked microtubules. (B) Schematic diagram showing the different functional domains of full-length CENP-E including the C terminus of CENP-E used in this study. (C) Representative images of HeLa cells transiently transfected with GST-GFP-CENP-E_2605-2701_ and immunostained with PRC1 and CENP-C, scale bar 10μm. (D) Sequence alignment of the C terminus of human CENP-E with 8 mammalian and Xenopus laevis CENP-E sequences. Amino acid numbering is relative to the human CENP-E sequence. The two PRC1 putative motifs ΦΦ are highlighted with stars (***). The following negatively charged amino acid is also highlighted. Phosphorylated residues following the CDK and Aurora kinase consensus sites are marked with an orange and blue circle respectively. Green circles represent sites that are phosphorylated but do not fit a kinase consensus site. The sequences were aligned using the program Clustal Omega (EBI) and formatted with ESPRIPT (Gouet et al., 1999). (E) Top, Representative images of live HeLa cells in cytokinesis transiently transfected with GFP-CENP-E_2605-2701_ and mutants (green), incubated with SiR-tubulin (red). Scale bar, 10μm. Bottom, linescans showing the fluorescence intensity average and standard error (SE) for the GFP-CENP-E_2605-2701_ and mutants and tubulin across the cell midbody. For GFP-CENP-E_2605-2701_, n=11 and for the GFP-CENP-E_2605-2701_ mutants FDN_2661_LTT, YF_2660_AA, FF_2644_AA, n= 15, 16 and 25 respectively. Experiments were repeated >3 times. (F) Graph showing the quantification of GFP peak fluorescence intensity for cells transfected with GFP-CENP-E_2605-2701_ constructs. Mean and peak intensities for indivual cells are represented for each mutant, ordinary One-way Anova test was performed to test significance.

PRC1 is a dimeric, non-motor, microtubule binding protein present on bundled spindle microtubules. It then is enriched on the central spindle in anaphase and in the midbody in telophase (Jiang et al., 1998; Mollinari et al., 2002). PRC1 preferentially binds to antiparallel overlapping microtubules (Bieling et al., 2010; Subramanian et al., 2010). Antiparallel microtubules compact to form a central spindle, to facilitate chromosome separation and specify the division plane. The timing of PRC1 recruitment coincides with the rapid dephosphorylation of the proteome during the metaphase to anaphase transition and is controlled by mitotic phosphorylation and dephosphorylation (Hu et al., 2012; Mollinari et al., 2002). Several motors — the Kinesin-4 Kif4A, Kif14A, MKLP1/Kif23 and CENP-E — are recruited to overlapping-microtubules in a PRC1-dependent manner in anaphase, and interact with PRC1 in cell extracts (Douglas and Mishima, 2010; Glotzer, 2009; Gruneberg et al., 2006; Hornick et al., 2010; Kurasawa et al., 2004).

Association of PRC1 with motor proteins controls the organization and the length of microtubule overlaps in the central spindle (Gruneberg et al., 2006; Kurasawa et al., 2004; Lee et al., 2015; Poser et al., 2019; Zhu and Jiang, 2005). Interestingly, Kif4A is bound to chromatin in early mitosis. It relocalizes in anaphase to the PRC1 marked-central spindle (Kurasawa et al., 2004; Wang and Adler, 1995). *In vitro*, PRC1 and Kif4A organize microtubules into bundles that resemble overlapping microtubule arrays in the central spindle. Kif4A enables motorized sliding of microtubules past each other in the presence of PRC1 (Bieling et al., 2010; Hannabuss et al., 2019; Subramanian et al., 2013).

In order to understand how PRC1 recruits and interacts with motors to assemble the central spindle and enable the timely completion of mitosis, we combine cell biology and structural approaches to dissect the molecular mechanism of the PRC1-motor interaction, focusing on how CENP-E is recruited to PRC1 in anaphase, for which little is known. We show that human CENP-E interacts directly with PRC1 using a bipartite ΦΦ motif at their C terminus, and this mechanism is also used by Kif4A to bind PRC1. We applied AlphaFold 2 to identify a region of PRC1 that is predicted to bind to CENP-E, and validated our predictions using site-directed mutagenesis. We used *in vitro* reconstitution and TIRF microscopy to show that full-length CENP-E slides PRC1-crosslinked microtubules past each other. Finally, using both biochemistry and cell biology approaches, we showed that the CENP-E:PRC1 interaction is spatially and temporally regulated by a phosphoswitch, which enables rapid relocalization of CENP-E from kinetochores to the central spindle during the metaphase to anaphase transition.

Our findings provide a framework for understanding how phosphorylation controls the spatial and temporal activities of kinesin motors using CENP-E as a paradigm, to enable completion of mitosis.

## RESULTS

### The C terminus of CENP-E co-localizes with PRC1 throughout mitosis

CENP-E localizes to the central spindle and midbody in anaphase and telophase (Kurasawa et al., 2004). Previous work indicated that this process requires the C terminus of CENP-E and is dependent on PRC1 (Fig 1B)(Ohashi et al., 2016). We hypothesized that the function of CENP-E in anaphase is independent of its kinetochore function. We have previously mapped the kinetochore-targeting domain (a.a. 2055-2608) (Chan et al., 1998; Legal et al., 2020). Thus we hypothesized the region of CENP-E that targets to the central spindle and PRC1 would be the region C-terminal to the kinetochore-targeting domain.

In order to mimic full-length CENP-E which is homodimeric, we dimerized CENP-E_2605-2701_ by fusing a GST-tag N-terminal to CENP-E_2605-2701_, as previously reported (Legal et al., 2020). To test whether the C-terminal fragment of CENP-E_2605-2701_ localized to overlapping microtubules in the central spindle during anaphase, and to the midbody during telophase when PRC1 is present, we imaged GST-GFP-CENP-E_2605-2701_ in mitosis. We found GST-GFP-CENP-E_2605-2701_ weakly co-localized with PRC1 to the centre of the metaphase spindle, and more strongly localized to PRC1-crosslinked microtubule bundles on the central spindle and midbody (Fig 1C). This is similar to the localization of full-length CENP-E in anaphase and telophase (Kurasawa et al., 2004).

### A Conserved CENP-E motif is required for PRC-1 binding

An alignment of full-length CENP-E from 9 mammalian species and *Xenopus laevis* revealed high levels of sequence conservation in the last 100 amino acids of CENP-E, C-terminal to the kinetochore-targeting domain (Fig 1B, D). This C-terminal domain contains the sequence motif RYFDNSSL (amino acids 2659-2666), which was previously reported to be essential for localization of CENP-E to the midbody. The localization of the CENP-E C terminus is also dependent on PRC1 (Ohashi et al., 2016).

We mutated the strongly conserved residues FDN (F2661, D2662, N2663) or YF (Y2660, F2661) to LTT and AA respectively. We then imaged cells in mitosis expressing wild type or mutant GFP-CENP-E_2605-2701_ using live-cell imaging to preserve dynamic interactions. Microtubules were stained with the SiR-tubulin dye, well suited for live-cell imaging. Both GFP-CENP-E_2605-2701_ mutants failed to localize to the midbody (Fig 1E). Interestingly, we also observed a similar, highly conserved (F or L/ F or L/ D) motif upstream, at position 2644-2646 (Fig 1D). To test the contribution of these two motifs to CENP-E recruitment to the central spindle and the midbody, we generated a series of mutations in CENP-E, that had altered motifs (either one motif, or the other, or both were mutagenized – alone or in tandem, Table 1).

**Table 1:** constructs generated in this study.

| Construct | Vector | Expression |
| --- | --- | --- |
| GFP-CENP-E 2605-2701 | pBabe blasticidin | human |
| MBP-CENP-E 2605-2701 | pMALC2X | bacteria |
| MBP-CENP-E 2605-2701 S2639D S2654D | pMALC2X | bacteria |
| MBP-CENP-E 2605-2701 F2661L, D2662T, N2663T | pMALC2X | bacteria |
| GFP-CENP-E 2605-2701 S2639D S2654D | pBabe blasticidin | human |
| GFP-CENP-E 2605-2701 F2661L, D2662T, N2663T | pBabe blasticidin | human |
| GFP-CENP-E 2605-2701 S2639A S2654A | pBabe blasticidin | human |
| GFP-CENP-E 2605-2701 Y2660A F2661A | pBabe blasticidin | human |
| GFP-CENP-E 2605-2701 F2644A F2645A | pBabe blasticidin | human |
| GST-CENP-E 2605-2701 | pGEX6p1 | bacteria |
| GST-CENP-E 2605-2701 S2639D S2654D | pGEX6p1 | bacteria |
| GST-CENP-E 2605-2701 S2639A S2654A | pGEX6p1 | bacteria |
| GFP-GST-CENP-E 2605-2701 | pBabe blasticidin | human |
| GFP-GST-CENP-E 2605-2701 S2639D S2654D | pBabe blasticidin | human |
| GFP-GST-CENP-E 2605-2701 S2639A S2654A | pBabe blasticidin | human |
| GST-CENP-E 2605-2701 F2644A F2645A Y2660A F2661A | pGEX6p1 | bacteria |
| GST-CENP-E 2605-2701 Y2660A F2661A | pGEX6p1 | bacteria |
| GST-CENP-E 2605-2701 F2644A F2645A | pGEX6p1 | bacteria |
| GST-CENP-E 2605-2701 F2661L, D2662T, N2663T | pGEX6p1 | bacteria |
| GFP-GST-CENP-E 2605-2701 F2661L, D2662T, N2663T | pBabe blasticidin | human |
| GST-CENP-E 2605-2701 S2639D S2647D S2469D S2651D S2654D S2664D | pGEX6p1 | bacteria |
| GST-CENP-E 2605-2701 S2639A S2647A S2469A S2651A S2654A S2664A | pGEX6p1 | bacteria |
| GST-CENP-E 2639-2671 | pGEX6p1 | bacteria |
| GFP-GST-CENP-E 2605-2701 S2639D S2647D S2469D S2651D S2654D S2664D | pBabe blasticidin | human |
| GFP-GST-CENP-E 2605-2701 S2639A S2647A S2469A S2651A S2654A S2664A | pBabe blasticidin | human |
| GST-Kif4A 1133-1165 | pGEX6p1 | bacteria |
| GST-Kif4A 1133-1232 | pGEX6p1 | bacteria |
| GFP-GST-Kif4a 1133-1232 | pBabe blasticidin | human |
| GFP-PRC1 guide RNA resistant | pBabe blasticidin | human |
| GFP-PRC1 3A M54A E57A E58A | pBabe blasticidin | human |
| GFP-PRC1 3A M54A E57A E58A guide RNA resistant | pBabe blasticidin | human |

Mutation _2644_FF_2645_ to AA caused a significant reduction in the localization of CENP-E to the midbody, whereas mutation of _2660_YF_2661_ to AA completely abolished CENP-E localization to the midbody (Fig 1E, F). We concluded that the second motif present in CENP-E, _2660_YF_2661,_ is essential for midbody localization, and that the _2644_FF_2645_ motif also contributes to targeting to the midbody. Taken together, our data reveal that the hydrophobic motif (Y or F), which we describe as ΦΦ motif, is essential for CENP-E recruitment to overlapping microtubules, and to PRC1 *in vivo*.

### CENP-E binds PRC1 *in vitro*

Next, we tested whether CENP-E and PRC1 interact *in vitro*. We expressed the C terminus of CENP-E as a monomeric MBP fusion, named MBP-CENP-E_2605-2701_ and a PRC1_1-168_ fragment, previously reported to bind to Kif4A (Subramanian et al., 2013) (Table 1). After mixing them together in an equimolar ratio and carrying out size-exclusion chromatography and SDS-PAGE analysis, we observed co-migration, indicating that these proteins assemble into a complex in solution (Fig 2A). We hypothesized that disruption of the second motif, which is essential for CENP-E recruitment to the central spindle, might abrogate the interaction of CENP-E with PRC1. In order to test this, we purified MBP-CENP-E_2605-2701_ mutant FDN_2661_LTT. Size-exclusion chromatography of MBP-CENP-E_2605-2701_ FDN_2661_LTT mixed with PRC1_1-168_ in a 1:1 molar ratio followed by SDS-PAGE analysis revealed that the two proteins no longer interacted. This confirms that the region flanking the second ΦΦ motif is essential for PRC1-CENP-E to interact *in vitro* (Fig 2B).

**Figure 2:**
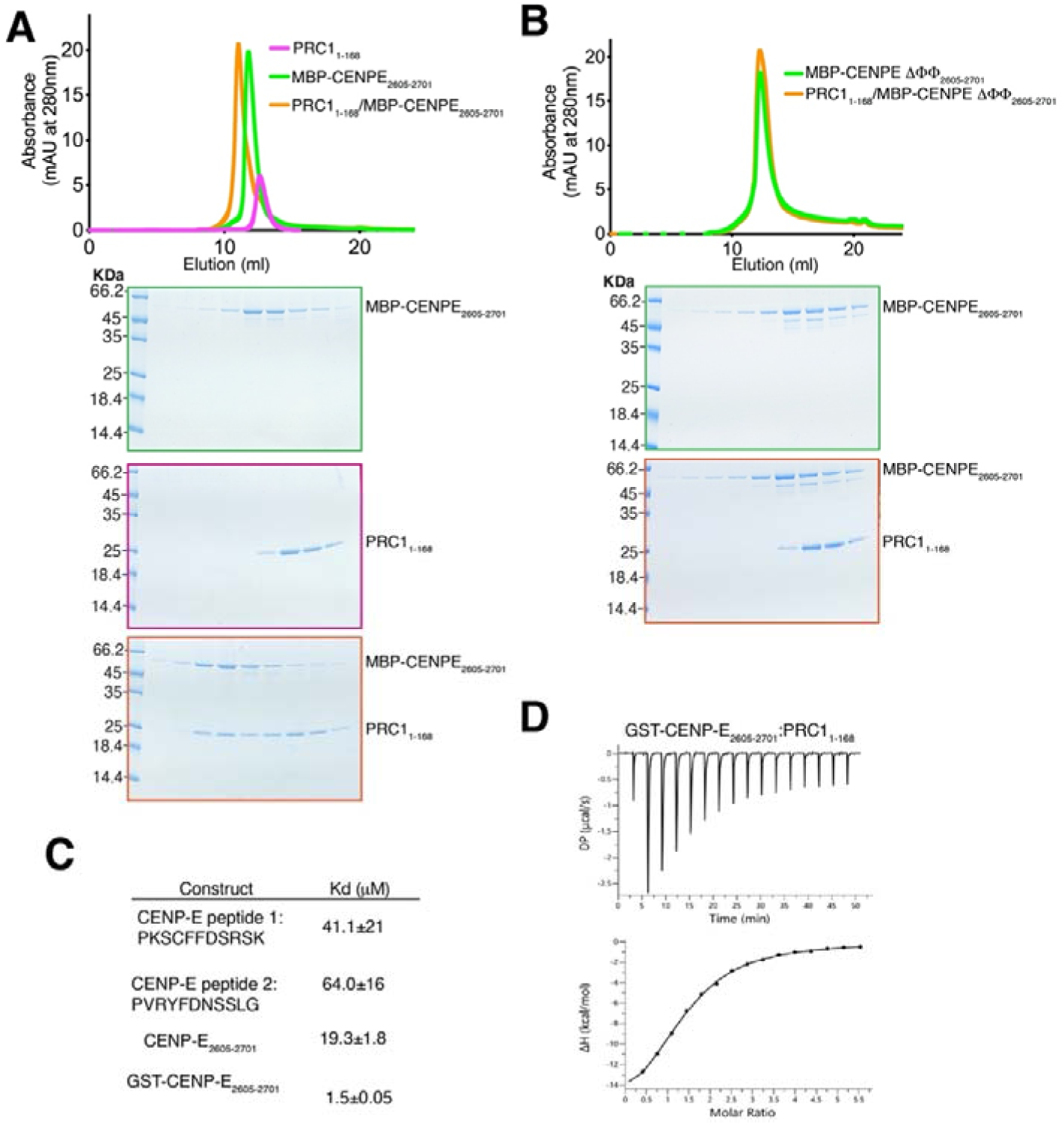
CENP-E interact with PRC1 through a kinesin ΦΦ motif. (A) Top. Size-exclusion chromatography elution profile of MBP-CENP-E_2605-2701_ (green), PRC1_1-168_ (pink) and MBP-CENP-E_2605-2701_/PRC1_1-168_ (orange). Bottom, coomassie-stained gel showing the size-exclusion chromatography profile of PRC1_1-168_ (pink), MBP-CENP-E_2605-2701_ (green) and MBP-CENP-E_2605-2701_/PRC1_1-168_ (orange). A shift in the elution volume was only seen in the presence of both CENP-E and PRC1. (B) Top. Size-exclusion chromatography elution profile of MBP-CENP-E_2605-2701_ ΔΦΦ (green) and MBP-CENP-E_2605-2701_ ΔΦΦ /PRC1_1-168_ (orange). Bottom, coomassie-stained gel showing the size-exclusion chromatography profile of MBP-CENP-E_2605-2701_ ΔΦΦ (green) and MBP-CENP-E_2605-2701_ ΔΦΦ /PRC1_1-168_ (orange). No shift in the elution profile was observed. (C) Table summarizing the isothermal titration calorimetry (ITC) results measuring the dissociation constant Kd for the PRC1_1-168_/CENP-E C-terminus interaction, using various C-terminal peptides. (D) Characterization by ITC of the PRC1_1-168_/GST-CENP-E_2605-2701_ interaction. Bottom. Top DP is the differential power and ΔH is the enthalpy.

### CENP-E binds PRC1 with high affinity

In cell extracts, PRC1 interacts with Kif4A, Kif14, MKLP1 and CENP-E (Kurasawa et al., 2004). The PRC1-Kif4A interaction has been reconstituted *in vitro* (Bieling et al., 2010; Subramanian et al., 2013) but the mechanism underlying the interaction of PRC1 with kinesin motors has not been reported. Hence, we sought to delineate the mechanism by which PRC1 binds to CENP-E by quantifying the affinity of CENP-E for PRC1. We used the previously characterized PRC1_1-168_ fragment which binds Kif4A (Subramanian et al., 2013). First, we measured the affinity of the CENP-E peptides containing the two ΦΦ motifs for PRC1 separately, to understand the contributions of these two sites. Isothermal calorimetry (ITC) measurement of CENP-E peptide 1, containing the first ΦΦ motif (PKSC_2644_FF_2645_DSRSK), and peptide 2, containing the second ΦΦ motif (PVR_2660_YF_2661_PNSSLG) affinity for PRC1_1-168_ revealed a very weak affinity of each peptide for PRC1 (Fig 2C, Fig S1A, B). Next we determined the affinity of the 98 amino acid C terminal fragment of CENP-E (CENP-E_2605-2701_), which contains both ΦΦ motifs in tandem in their native CENP-E sequence (Fig S1C) with PRC1_1-168_. We measured an affinity of 19.3 μM (Fig 2C). The C-terminal domain of CENP-E is predicted to be disordered using AlphaFold2 (Jumper et al., 2021), so binding to PRC1 would likely involve a large entropic penalty, associated with a reduction in conformational flexibility in the protein upon binding to PRC1. Overall, this interaction is relatively weak. However, CENP-E and PRC1 are both dimers *in vivo*. So, in order to more closely represent the *in vivo* interaction, we purified GST-CENP-E_2605-2701_ which is dimeric and measured the affinity of GST-CENP-E_2605-2701_ for PRC1_1-168_ using ITC (Fig S1D). Dimerization led to a 12-fold increase in affinity between GST-CENP-E_2605-2701_ for PRC1_1-168_ to 1.5 μM, similar to the binding affinity reported for the Kif4A:PRC1 measured using binding assays (Subramanian et al., 2013). Overall these data indicate multiple ΦΦ motifs increase the affinity of the CENP-E:PRC1 interaction through an avidity effect.

### Kif4A requires a bipartite ΦΦ motif for PRC1 binding

Kif4A also contains a phenylalanine ΦΦ motif (F1154, F1155) essential for targeting to the central spindle and PRC1 binding (Poser et al., 2019). It is reminiscent of the CENP-E motif (Fig 3A), although no aspartate follows the ΦΦ motif. We expressed the Kif4A 32 amino acid region containing the ΦΦ motif as a GST fusion (GST-Kif4A_1133-1165_)(Table 1). Unlike the C-terminus of CENP-E, GST-Kif4A_1133-1165_ did not have any affinity for PRC1, as measured by ITC (Fig S1E). *In vivo*, GFP-GST-Kif4A_1133-1165_ did not localize to overlapping microtubules (Fig 3B). We then searched for a second motif that could increase Kif4A binding to PRC1, similarly to CENP-E. There is a second ΦΦ motif in Kif4A (F1220, F1221) upstream of the published PRC1-binding region (F1154, F1155) (Fig 3B). We hypothesized that this second motif might contribute to the PRC1-Kif4A interaction, and that the ΦΦ motif (F1154, F1155) is essential but not sufficient for PRC1 binding, in common with CENP-E (Fig S1). To test whether Kif4A and CENP-E bind PRC1 using a similar mechanism, we expressed a dimeric fragment of Kif4A that contains both ΦΦ motifs, GFP-GST-Kif4A_1133-1232,_ and showed that it localizes to overlapping microtubules (Fig 3B). *In vitro*, GST-Kif4A_1133-1232_ and PRC1 co-eluted as a complex using SEC (Fig 3C, D). Together, these data indicate that both while the previously reported Kif4a ΦΦ motif is necessary to bind PRC1 (Poser et al., 2019), it is not sufficient. Similarly to CENP-E, Kif4a uses a bipartite motif to stably bind PRC1.

**Figure 3:**
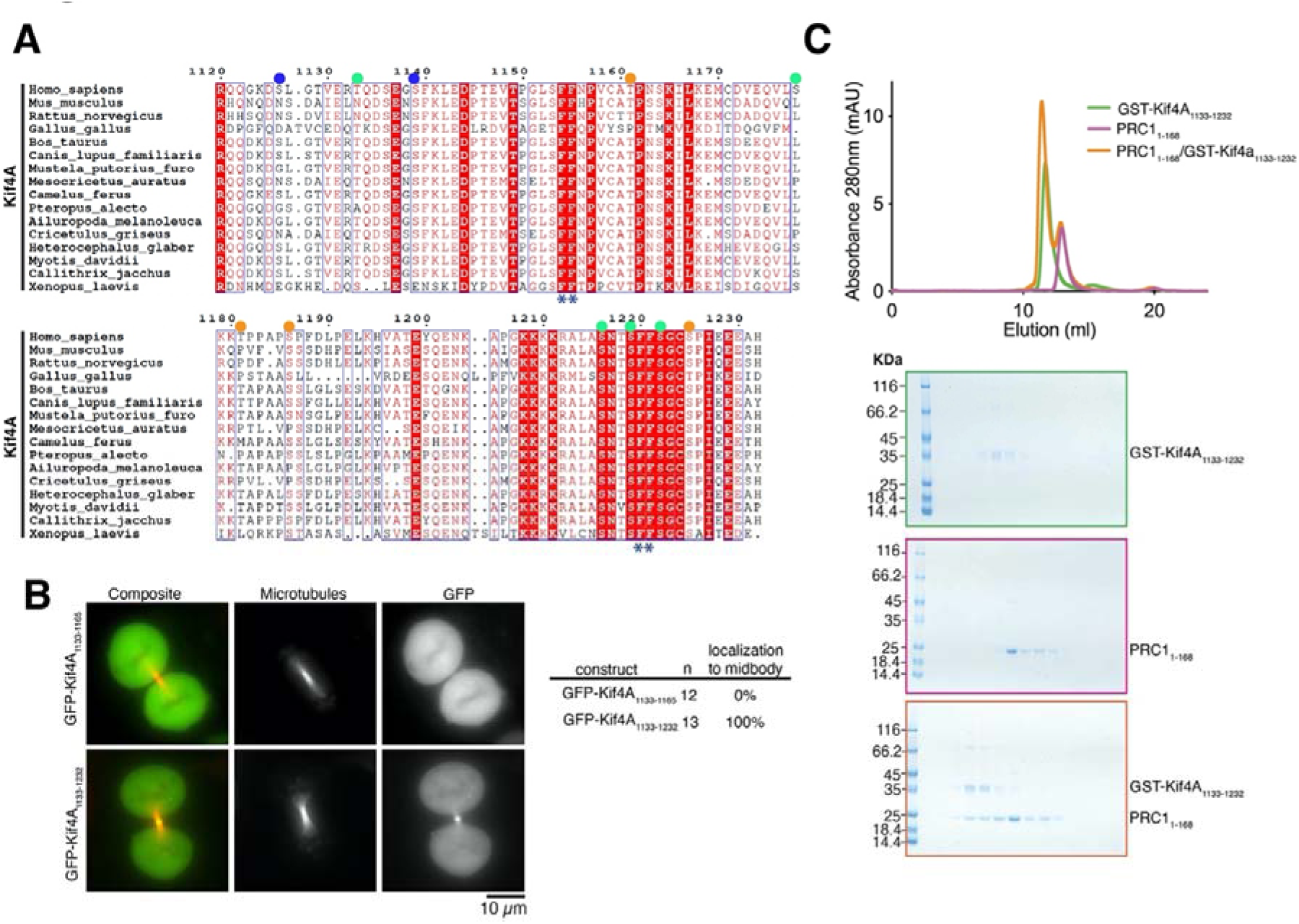
Kif4a binds to PRC1 using a bipartite ΦΦ motif. (A) Sequence alignment of the C terminus of human Kif4a with Kif4a of other metazoans. Amino acid numbering is relative to the human Kif4a sequence. The two PRC1 putative motifs ΦΦ are highlighted with stars (*). Published phosphorylated residues are marked. Those following the CDK and Aurora kinase consensus sites are marked with an orange and blue circle respectively. Green circles represent sites that are phosphorylated but do not fit a kinase consensus site. The sequences were aligned using the program Clustal Omega (EBI) and formatted with ESPRIPT (Gouet et al., 1999). (B) Representative images of live HeLa cells in mitosis transiently transfected with either GFP-Kif4a_1133-1165_ or GFP-Kif4a_1133-1232_ incubated with SiR-Tubulin. Scale bar, 10μm. Quantification of GFP-Kif4a_1133-1165_ (n=12) or GFP-Kif4a_1133-1232_ (n=13) localization to microtubules at the midbody. (C) A shift in the elution volume was only seen in the presence of both Kif4a and PRC1. Size-exclusion chromatography elution profile of GST-Kif4a_1133-1232_ (green) and PRC1_1-168_ (pink), or together (orange). Bottom, coomassie-stained gel showing the size-exclusion chromatography profile of GST-Kif4a_1133-1232_ (green), PRC1_1-168_(pink) and GST-Kif4a_1133-1232_/PRC1_1-168_ (orange).

### A CENP-E-PRC1 complex slides microtubules

CENP-E has been proposed to slide microtubules in mitosis using its non-motor microtubule binding domain (Steblyanko et al., 2020). It is also possible that CENP-E slides antiparallel microtubules that are crosslinked by PRC1, similarly to Kif4A (Bieling et al., 2010; Subramanian et al., 2013). To distinguish between these two models, we carried out an *in vitro* reconstitution experiment (Fig 4). We previously reconstituted motility of both truncated and full-length CENP-E *in vitro* (Craske et al., 2022). The challenge in analyzing the contribution of CENP-E to microtubule sliding is that about 10% purified full-length CENP-E is motile, with the long coiled-coil stalk interfering with its activity (Fig S2A)(Craske et al., 2022). In order to determine whether CENP-E slides microtubules alone, or only slides those crosslinked by PRC1, we first analyzed whether CENP-E was recruited to microtubules crosslinked with PRC1, or to PRC1 directly. As full-length CENP-E is challenging to work with owing to its instability, we purified a ‘minimal PRC1-binding CENP-E construct’, GST-CENP-E_2639-2671_ (Table S1)(Fig S2A) and chemically labelled this protein with an Alexa Fluor-647 dye. To test if PRC1 is able to recruit the CENP-E C terminus to microtubules, polymerized GMPCPP-stabilized rhodamine microtubules were mixed with _647_GST-CENP-E_2639-2671_ alone, GFP-PRC1 alone or both _647_GST-CENP-E_2639-2671_ and GFP-PRC1. These samples were then added to silanised coverslips that were coated with anti-tubulin antibodies (Fig 4 A-C). When we added GFP-PRC1 to GMPCCP-stabilized rhodamine microtubules in a flow chamber, GFP-PRC1 decorated the length of the microtubule but was preferentially recruited to overlapping regions between two or more microtubules (Fig 4B). In the presence of GFP-PRC1, _647_GST-CENP-E_2639-2671_ bound specifically to PRC1 at overlapping microtubules. These results indicate that the C terminus of CENP-E specifically binds to PRC1 rather than to microtubules (Fig 4C). _647_GST-CENP-E_2639-2671_ also recognizes and stains endogenous PRC1 in cells (Fig 4D).

**Figure 4:**
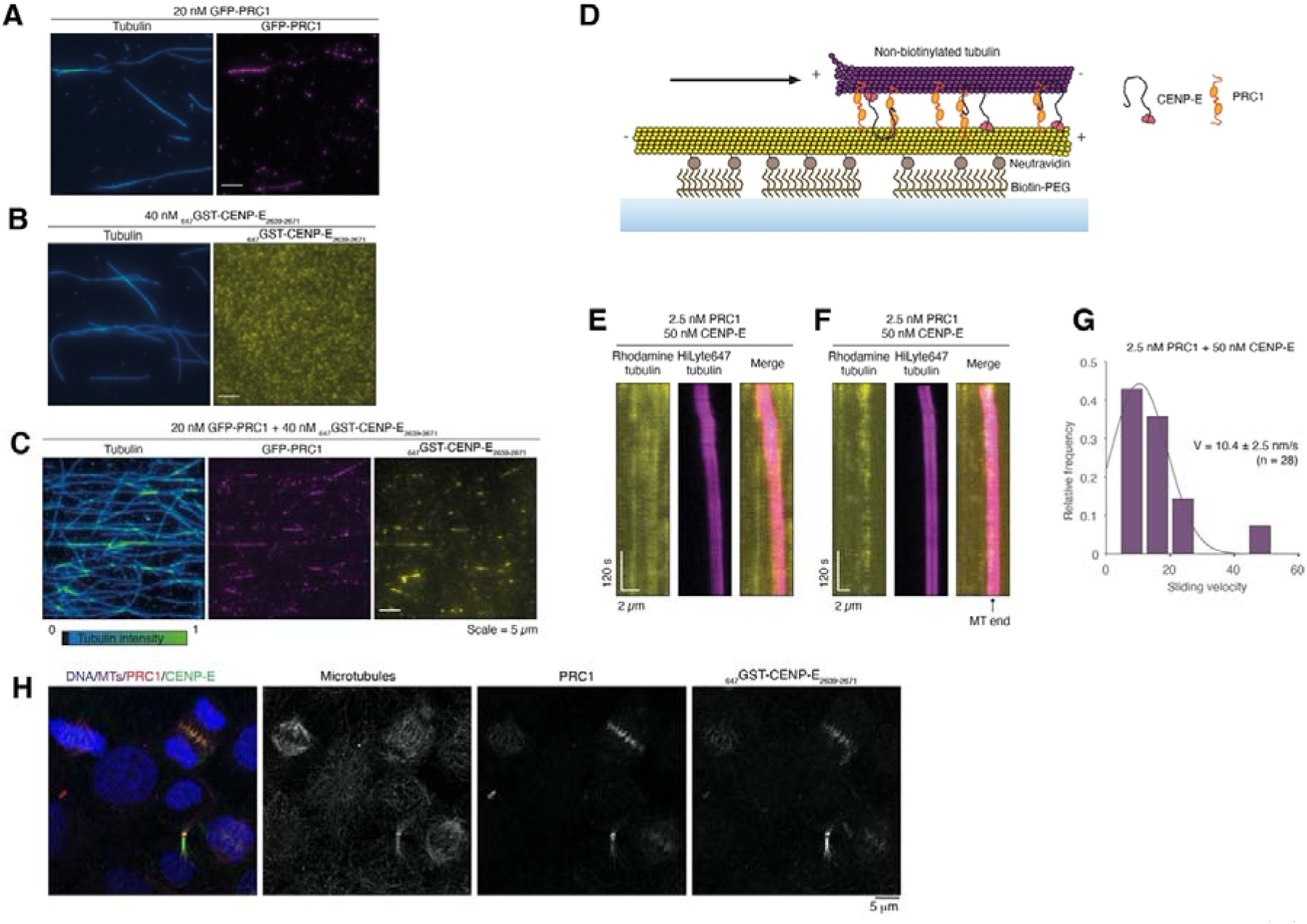
CENP-E slides anti-parallel microtubules in the presence of PRC1. (A) Representative images of GFP-PRC1 (magenta) mixed with rhodamine-microtubules. Fire blue-green intensity LUT used to show tubulin intensity and microtubule overlaps. (B) Representative images of _647_GST-CENP-E_2639-2671_ (yellow) mixed with rhodamine-microtubules. (C) Representative images of GFP-PRC1 and GST-CENP-E_2639-2671_ mixed with rhodamine-microtubules. (D) Schematic representation of sliding in reconstituted system. (E) Representative kymograph showing microtubule-microtubule sliding in the presence of 2.5 nM PRC1 and 50 nM full-length CENP-E. (F) Example kymograph showing a free microtubule sliding until reaching the end of the immobilized microtubule where it slows down to a stall. (G) Graph showing the quantification of microtubule-microtubule sliding velocity exhibited by free microtubules transported in the presence of 2.5 nM PRC1 and 50 nM full-length CENP-E, n=28. (H) Representative immunofluorescence images of HeLa cells stained for DNA, microtubules, PRC1 and with Alexa647-labelled GST-CENP-E_2639-2671_ showing it also recognizes PRC1 in cells. Experiments were repeated >3 times.

Next, we analyzed whether full-length human CENP-E could slide microtubules apart in the presence of PRC1 *in vitro*. We incubated surface immobilized microtubules with full-length human PRC1, to allow coating of the microtubule lattice with PRC1, then added GMPCPP-stabilized rhodamine microtubules, which led to microtubule bundling (Fig 4E). When 2.5 nM PRC1 alone was added, pairs of overlapping microtubules formed (Fig S2B). Overlaps remained constant over time, and no sliding of microtubules was observed throughout the experiment lasting 20 minutes (Fig S2C). In contrast, we did not observe overlapping microtubule pairs when we added 50 nM full-length CENP-E, ATP and microtubules (Fig S2B). When we added 50 nM CENP-E and 2.5 nM PRC1, free microtubules were crosslinked and transported unidirectionally along the coverslip-immobilized microtubules (Fig 4F and G, S2B, movie 1). CENP-E-driven microtubule sliding was slow, with an average velocity of 10.4 ± 2.5 nm/s (Fig 4H). This sliding velocity is comparable to that of Kif4A, around 11nm/s in the presence of 1nM PRC1)(Wijeratne and Subramanian, 2018).

Together, these data suggest that a CENP-E-PRC1 complex is able to slide antiparallel microtubules relative to each other. The sliding velocity may be regulated by frictional forces that are either generated by the accumulation over time of PRC1 on cross-linked microtubules, similar to Kif4A/PRC1 sliding (Wijeratne and Subramanian, 2018), or by the fraction of inactive or paused microtubule-bound CENP-E motors that can still bind to PRC1 (Craske et al., 2022).

### Phosphorylation of CENP-E controls its PRC1-microtubule binding activity

CENP-E is only recruited to overlapping antiparallel microtubules, crosslinked by PRC1 in anaphase. Before anaphase, CENP-E is primarily localized to unattached kinetochores in prometaphase, and remains localized to kinetochores in smaller amounts in metaphase. We hypothesized that the interaction between PRC1 and CENP-E might be regulated by post-translational modifications to enable rapid temporal and spatial relocalization of CENP-E from kinetochores to PRC1-bound microtubules in the central spindle. Of note, mitotic kinase activity is high in prometaphase, contributed by CDK, Aurora, Mps1 and Plk1 kinases. In particular, CDK activity drops dramatically during the metaphase to anaphase transition. Multiple phosphoproteomic studies have previously reported that the C terminus of CENP-E is phosphorylated in mitosis and identified the phosphorylated residues *in vivo* (Dephoure et al., 2008; Kettenbach et al., 2011; Malik et al., 2009; Nousiainen et al., 2006; Santamaria et al., 2011; Sharma et al., 2014). We noted 6 of these phosphorylated residues were close to the PRC1 binding motif (Fig 1D). Two serines phosphorylated at position 2639 and 2654 fit the CDK consensus site (S/T-P) and a serine 2651 phosphorylated by the Aurora kinases close to the FF motifs were reported multiple times (Kettenbach et al., 2011; Nousiainen et al., 2006; Santamaria et al., 2011; Sharma et al., 2014)(Fig 1D). S2647, S2649 and S2664 were also reported as phosphosites (Sharma et al., 2014).

In order to test whether phosphorylation of the C terminus of CENP-E affects its interaction with PRC1, we generated phosphomimetic (amino acid substitutions that mimic a phosphorylated version of the amino acid) mutants of GST-CENP-E_2605-2701_. We mutated S2639 and S2654 to generate GST-CENP-E_2605-2701_ 2SD (mimicking 2 phosphorylated amino acids), and S2639, S2647, S2649, S2651, S2654 and S2664 for GST-CENP-E_2605-2701_ 6SD (mimicking 6 phosphorylated amino acids), and measured their affinity for PRC1 using ITC. There was a small decrease in affinity of GST-CENP-E_2605-2701_ 2SD, (Kd increased from 1.5 μM for control versus 2.4 μM for 2SD), and GST-CENP-E_2605-2701_ 6SD displayed no binding to PRC1 (Fig S1E-G). Further phosphorylation of CENP-E could also contribute to reducing the PRC1:CENP-E interaction *in vivo*.

Next, we analyzed how phosphorylation of the C terminus of CENP-E affected association with PRC1 at overlapping microtubules in cells (Fig 5). We examined the localization of CENP-E C terminus in metaphase (Fig 5A). GFP-CENP-E_2605-2701_ was mostly cytoplasmic, but dimeric GFP-GST-CENP-E_2605-2701_, (as full-length CENP-E functions as a dimer *in vivo*), was enriched on interpolar overlapping microtubules, close to the chromosomes. It was not observed uniformly on microtubules, indicating GST-CENP-E_2605-2701_ is unlikely to bind microtubules directly (Fig 5A). GFP-GST-CENP-E_2605-2701_ 2SD localized weakly to interpolar microtubules, but GFP-CENP-E_2605-2701_ 2SD and GFP-GST-CENP-E_2605-2701_ 6SD did not associate with microtubules at all. We also frequently observed GFP-CENP-E_2605-2701_ 2SA, albeit in small amounts (weak fluorescence) on interpolar microtubules in metaphase. GFP-GST-CENP-E_2605-2701_ 2SA is dimeric and was observed on interpolar microtubules (Fig 5A). GFP-GST-CENP-E_2605-2701_ 6SA was also enriched on overlapping microtubules in the mitotic spindle.

**Figure 5:**
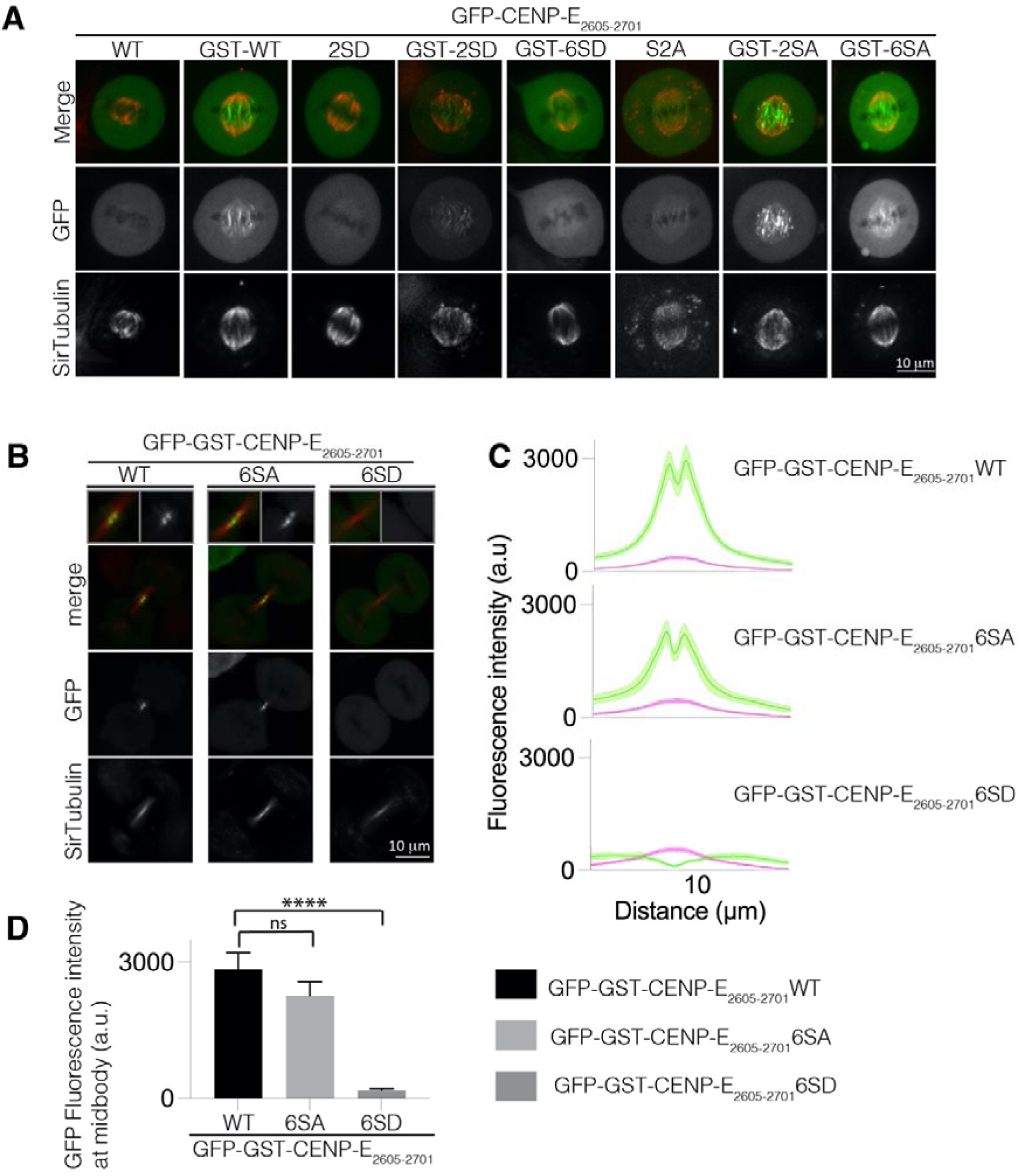
Phosphorylation state of CENP-E C terminus regulates its localization to overlapping microtubules. (A) Live-cell imaging of metaphase spindles in HeLa cells transfected wild type (n=27), phosphomimetic (n=20) and non-phosphorylatable (n=21) mutants of GFP-CENP-E_2605-2701_ (monomeric) and GFP-GST-CENP-E_2605-2701_ (dimeric) and stained for tubulin using SiR-Tubulin. Scalebar, 10 μm. (B) Live-cell imaging of the midbody in HeLa cells transfected with wild type, phosphomimetic and non-phosphorylatable mutants of GFP-GST-CENP-E_2605-2701_ and stained for tubulin. Scalebar, 10 μm. (C) Linescans showing the mean fluorescence intensity and standard error (SE) for the GFP-CENP-E_2605-2701 wild type_ constructs and mutants and tubulin across the cell midbody. Experiments were repeated 2-3 times. (D) Bar graph showing mean and standard error for GFP fluorescence intensity at peak fluorescence for GFP-GST-CENP-E_2605-2701_ and mutants at 9.7μm, quantified in C. Asterisks indicate ordinary One-way Anova test significance value. ****P<0.0001.

Next, we examined the localization of GFP-GST-CENP-E_2605-2701_ at the midbody in telophase. GFP-GST-CENP-E_2605-2701_ and GFP-GST-CENP-E_2605-2701_ 6SA were present at the midbody but GFP-GST-CENP-E_2605-2701_ 6SD did not associate with the midbody, similar to our observations with the GFP-CENP-E_2605-2701_ YF2660AA mutant.

We surmise that phosphorylation of the CENP-E C terminus prevents association with PRC1. Taken together these data reveal that phosphorylation of the C terminus of CENP-E inhibits recruitment to PRC1 at overlapping microtubules in early mitosis, by reducing the affinity of CENP-E for PRC1. The phosphorylation state of CENP-E during mitosis therefore regulates its interactions, both spatially and temporally, to enable CENP-E to associate with the outer corona of kinetochores in early mitosis, where it mediates chromosome capture and alignment, and then to associate with PRC1 later in mitosis.

### Structural features of CENP-E-PRC1 interactions

We used AlphaFold2 to predict how CENP-E might interact with PRC1, using CENP-E_2605-2701_ and PRC1_1-168_ dimers as inputs for our analysis (Jumper et al., 2021; Mirdita et al., 2022). AlphaFold2 predicted that CENP-E could interact with the rod and dimerization interface of PRC1 with high confidence, and identified _2660_YFD_2661_ in CENP-E as important for that interaction (Fig S3A). PRC1 is dimeric, with the dimerization domains and the rod fold organized around a two-fold symmetry, antiparallel to each other. The ΦΦ binding sites on PRC1 are in close proximity to each other. This could explain why two ΦΦ motifs from the same peptide can bind to PRC1 to increase motor affinity, such as CENP-E or KIFA4, for PRC1. Based on AlphaFold2 predicted structures of the CENP-E_2605-2701_:PRC1 complex we could identify several amino acids that might be involved in coordinating the ΦΦ motif: I25, W26, M54, E57, E58. In order to test these structural predictions, we generated a PRC1_1-168_ in which M54, E57 and E58 were all mutated to A, named PRC1_1-168_ 3A (Table S1). PRC1_1-168_ 3A was soluble and behaved similarly to PRC1_1-168_ in size-exclusion chromatography (Fig S3B). However PRC1_1-168_ 3A could not bind to MBP-CENP-E_2605-2701_ (Fig S3B).

Overall, these data indicate that kinesin motors bind the dimerization-rod domain of PRC1 using their bipartite ΦΦ motifs.

### PRC1-motor interactions are essential for cytokinesis

PRC1 has dual molecular functions: it crosslinks microtubules, and it associates with kinesin motors, such as CENP-E and Kif4A. These functions allow assembly of the central spindle and ensure the final steps of mitosis. In order to distinguish the contribution(s) of PRC1 to central spindle formation, which could occur either by crosslinking microtubules or by recruiting kinesin motors, we engineered cell lines expressing GFP-PRC1-WT, or GFP-PRC1-3A (which does not bind the ΦΦ motif in kinesin motors) (Fig 6A). Both cell lines were stable, indicating that these constructs did not likely have a dominant effect. Both cell lines also expressed a guide RNA that targets endogenous PRC1 constitutively, with Cas9 expressed using an inducible promoter, so that we could induce PRC1 knockout by addition of doxycycline (McKinley and Cheeseman, 2017), as an alternative to siRNA knockdown (Fig 6A).

**Figure 6:**
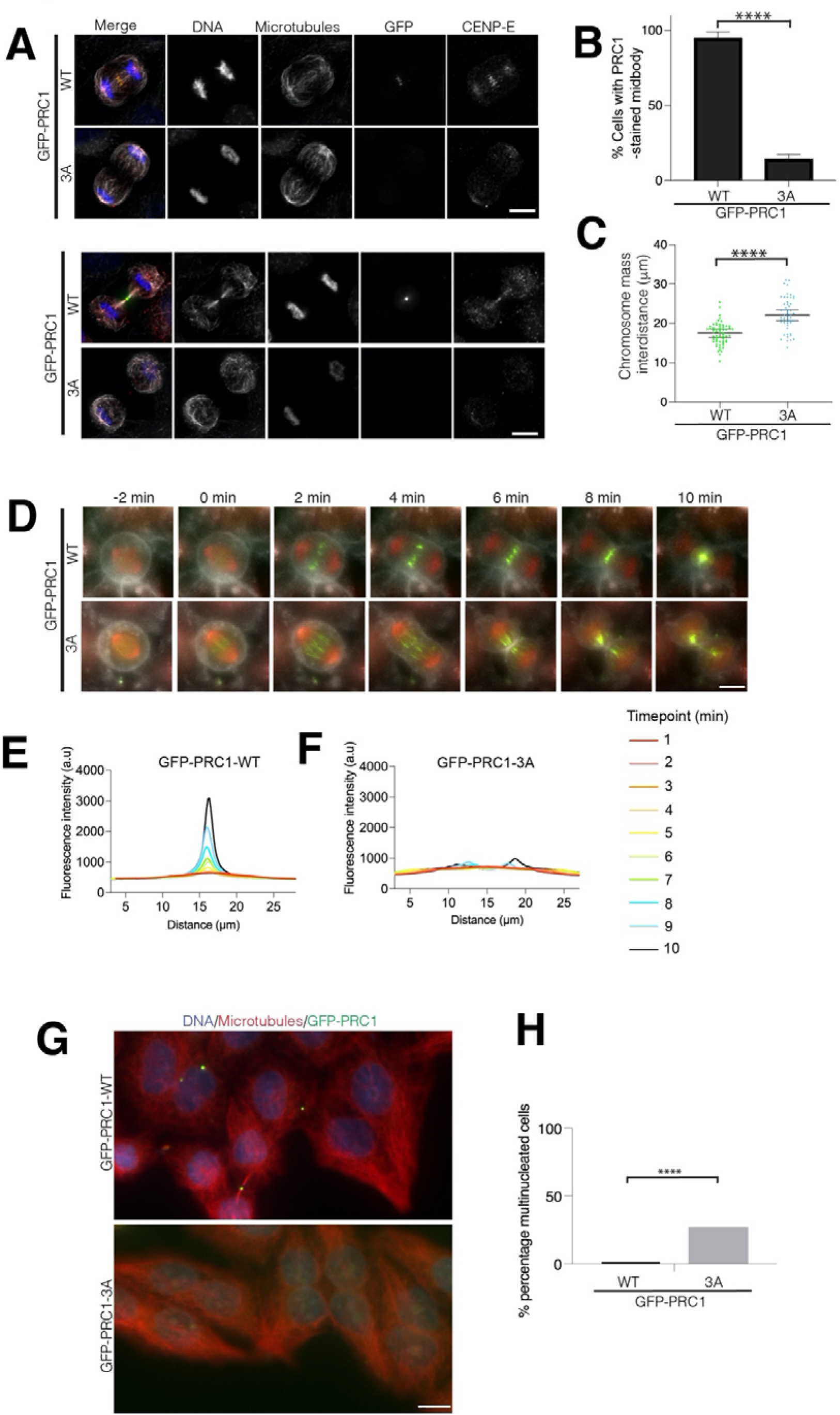
the interaction of PRC1 with motors is critical for the integrity of the central spindle. (A) Representative immunofluorescence images of HeLa cells expressing GFP-PRC1 wild type or 3A mutant and depleted for endogenous PRC1 using a PRC1 siRNA and a PRC1 sgRNA after doxycycline-induced Cas 9 expression. Microtubules, GFP and CENP-E are in white, green and red respectively. DNA is in blue. (B) Interchromosome distance in telophase in cells expressing GFP-PRC1 WT and GFP-PRC1 3A in the absence of endogenous PRC1 (n=60 and n=53 respectively, experiment repeated twice. (C) Graph showing the percentage of cells with a PRC1-localized midbody for cells treated as in (A). Asterisks indicate a Fisher’s exact test significance value. ****P<0.0001. (D) Representative live-cell images of HeLa cells expressing GFP-PRC1 wild type or 3A mutant and depleted for endogenous PRC1 using a PRC1 siRNA and a PRC1 sgRNA after doxycycline-induced Cas 9 expression. Microtubules (SiR-Tubulin) and DNA (SPY650) are shown in red. The cell membrane (CellMask) is highlighted in white and GFP-PRC1 is in green. Scalebar : 10 μm. (E-F) Graph showing the change in fluorescence intensity for DNA and microtubules along the longitudinal axis of the spindle from metaphase (red) to anaphase and cytokinesis (dark blue) for cells expressing GFP-PRC1 WT (E) or 3A (F) in the absence of endogenous PRC1. (G) Representative immunofluorescence images of cells expressing GFP-PRC1-WT or -3A after 72 hours siRNA depletion and doxycycline-induced knockout of endogenous PRC1. DNA, microtubules and GFP-PRC1/GFP-PRC1-3A are in blue, red and green respectively. Scalebar, 10μm. Experiments repeated 2-3 times. (H) Quantification of the number of cells mononucleated and multinucleated from experiment in (G). n=286 and 393 for cells expressing GFP-PRC1 WT and GFP-PRC1 3A respectively. Asterisks indicate a Fisher’s exact test significance value. ****P<0.0001.

We observed that GFP-PRC1-WT localizes to overlapping microtubules in the metaphase spindle, the central spindle and the midbody, as previously reported (Figure 6A, B)(Kajtez et al., 2016; Pamula et al., 2019; Subramanian et al., 2013). We found that GFP-PRC1-WT localization was slightly reduced when endogenous PRC1 was knocked down, most likely due to the N terminal GFP-tagging of PRC1, which places the tag close to the dimerization interface and the motor-binding interface. However, we observed that endogenous CENP-E was present at overlapping structures bound by GFP-PRC1-WT, indicating that GFP-PRC1-WT was able to interact with the ΦΦ motif of CENP-E and recruit CENP-E (Fig 6A). When endogenous PRC1 was knocked down, cells expressing GFP-PRC1-3A still progressed through mitosis and chromosome segregated in anaphase, meaning that checkpoint silencing must have taken place. In anaphase, the amount of GFP-PRC1-3A was reduced on the central part of the spindle compared with that observed for GFP-PRC1-WT (Fig 6A, B). The central spindle, which is usually marked by a high density of antiparallel microtubules, was absent or severely disrupted in cells expressing GFP-PRC1-3A and lacking endogenous PRC1. Very few microtubules were observed in between the two segregating half-spindles. CENP-E was not present in the central spindle in these cells (Fig 6A, B). Under these conditions, we observed hypersegregation of chromosomes in the presence of GFP-PRC1-3A, with the distance between the chromosome mass significantly greater than in cells expressing GFP-PRC1-WT (Fig 6A, C). This phenotype is similar to that seen in cells depleted for PRC1 (Pamula et al., 2019).

To better understand how PRC1-motor interactions affects chromosome segregation and cell division, we carried out time-lapse imaging on cells expressing GFP-PRC1-WT or GFP-PRC1-3A in the absence of endogenous PRC1, using dyes to demarcate cell membrane, DNA and microtubules. We observed that the speed of the chromosome and spindle pole mass to the daughter cells, marked by DNA and tubulin dyes respectively, was similar in cells expressing GFP-PRC1-3A compared with those expressing GFP-PRC1-WT (Fig 6E). We measured the accumulation of GFP-PRC1-WT on the central spindle over time (Fig 6F). GFP-PRC1-3A was weakly recruited to overlapping microtubules, unlike GFP-PRC1-WT, which accumulated in the central spindle throughout anaphase and telophase. However, there were less antiparallel microtubules and the fibres appear thicker, indicating an abnormal regulation of microtubule bundling in the presence of GFP-PRC1-3A (Fig 6D, F). The staining of GFP-PRC1-3A was more diffuse than for GFP-PRC1-WT, and the distribution of GFP-PRC1-3A was not constrained to the central spindle. Additionally, GFP-PRC1 marked bundles seemed to move away from each other and make thicker bundles, which were easier to distinguish during spindle elongation in anaphase. It is also possible the PRC1-marked bundles were severed or broken at the site of furrow ingression (Fig 6D).

PRC1 is essential for cytokinesis (Jiang et al., 1998; Mollinari et al., 2005). Next we analyzed whether the motor-recruitment properties of PRC1 contribute to cytokinesis. We depleted endogenous PRC1 in cells expressing GFP-PRC1-WT or GFP-PRC1-3A and imaged them after 72 hours. In cells expressing GFP-PRC1-3A, there was a significant increase in multinucleated cells, indicating them had failed cytokinesis and were polyploid. Overall, these results indicate the motor-recruitment properties of PRC1 are critical to ensure the completion of cytokinesis and support successful cell division. Future work will be needed to determine the contributions of individual PRC1-motor interactions to anaphase and cytokinesis.

## Discussion

In this manuscript, we report the mechanistic basis of PRC1-microtubule motor interactions, and recapitulate microtubule sliding by a CENP-E:PRC1 complex. We reveal the functional contribution of PRC1-interacting microtubule motors to spindle elongation in anaphase and the completion of chromosome segregation, identify the temporal and spatial features of these regulated interactions, and show that they are required for correct timing of cytokinesis.

In early mitosis, CENP-E is present at unattached kinetochores, and moves laterally-attached kinetochores along microtubules. The kinetochore-bound CENP-E has previously been proposed to slide microtubules by pushing microtubules relative to kinetochores and promote spindle flux (Steblyanko et al., 2020). CENP-E has also been proposed to slide spindle microtubules past each other (Risteski et al., 2021). Some motors, such as the Kinesin-14 dimeric motor HSET, use both its motor and non-motor microtubule binding domains to slide microtubules past each other (Braun et al., 2017; Cai et al., 2009). We show that CENP-E, unlike Kinesin-14 HSET, does not slide microtubules on its own *in vitro*.

Previous work on the CENP-E C-terminal tail included part of the kinetochore targeting domain and a His tag, which may have artificially promoted association with the negatively charged microtubule lattice (Gudimchuk et al., 2013). *In vitro* and *in vivo*, we did not observe any binding of the CENP-E C-terminal domain to microtubules in the absence of CENP-E interaction with PRC1 (Fig 4A). The unstructured C terminus of CENP-E is phosphorylated in metaphase (Dephoure et al., 2008). Therefore its affinity for microtubules in the context of the full-length motor is likely to be weak and non-specific. The data we present in this manuscript rule out microtubule sliding activity of CENP-E via its non-microtubule binding domain. Here we demonstrate that CENP-E promotes microtubule-microtubule sliding later in anaphase, and functions as a complex of CENP-E:PRC1, in a similar manner to Kif4A (Bieling et al., 2010; Subramanian et al., 2013)(Fig 2, 4).

The timing of the PRC1:motor interaction is important, because Kif4A and CENP-E are involved in chromosome organization and alignment in early mitosis (reviewed in (Craske et al., 2022; Samejima et al., 2012). PRC1 is phosphorylated by CDK1/cyclin B on T470 and T481, in the region important for microtubule binding, at the junction between the unstructured microtubule binding tail and the spectrin domain (Jiang et al., 1998). The microtubule bundling activity of PRC1 is downregulated by CDK1 phosphorylation (Mollinari et al., 2002). We show here that mitotic phosphorylation also regulates the PRC1:CENP-E interaction by phosphorylating CENP-E. Phosphorylation of CENP-E in prometaphase and metaphase ensures that CENP-E associates with kinetochores to promote their alignment. At this time CENP-E affinity for PRC1 is low. As chromosomes bi-orient and segregate, CENP-E is dephosphorylated, which increases its affinity for PRC1 and facilitates its recruitment to the central spindle. Interestingly, the two ΦΦ motifs in the KIF4A motor are also flanked by threonines and serines (Huttlin et al., 2010; Kettenbach et al., 2011; Nousiainen et al., 2006; Olsen et al., 2010)(Fig 3A). Thus mitotic phosphorylation of the C-terminus of Kif4A may regulate the KIF4A:PRC1 interaction temporally and spatially.

The motor-recruiting function of PRC1, via the ΦΦ binding site, is crucial to the correct completion of cytokinesis and chromosome segregation (Fig 6). When this interaction is abrogated, the chromosomes hypersegregate. Ultimately, cells fail cytokinesis and become multinucleated (Fig 6). It is possible that the MKLP2-PRC1 interaction is disrupted, if MKLP2 also uses a ΦΦ motif for PRC1 binding. This would prevent recruitment of the CPC (Chromosomal Passenger Complex) to the central spindle to complete cytokinesis (Adriaans et al., 2020; Gruneberg et al., 2004; Kitagawa et al., 2013; Serena et al., 2020).

During anaphase, we observe the central spindle starts to assemble, but the antiparallel microtubule bundles, marked by GFP-PRC1-3A, are reduced and no longer concentrated in the central spindle. GFP-PRC1-3A still has a strong preference for antiparallel microtubule fibres and crosslinks microtubules within the spindle overlap. However, GFP-PRC1-3A is not concentrated at the plus end of overlaps, highlighting that motors are essential for concentrating PRC1 and marking the central spindle and midbody (Hannabuss et al., 2019; Subramanian et al., 2013; Wijeratne and Subramanian, 2018). The PRC1-marked bundles also lose their coherent behavior within the spindle, and move away from each other. Recent work proposes that microtubule bundles in the central spindle are connected to each other (Carlini et al., 2022). Our data reveal that PRC1-interacting proteins that bind to the dimerization domain, contribute to interbundle stability by crosslinking different sets of microtubule bundles. This may reinforce their stiffness.

PRC1 is proposed to act as a break counteracting forces that drive spindle elongation (Janson et al., 2007).The forces generated by single PRC1 molecules on microtubules are low, in the 0.1 pN range (Forth et al., 2014). At higher density, PRC1-crosslinked microtubules can produce significant resistance during microtubule sliding, that scale with velocity of microtubule sliding, in the range 5-20 pN for sliding velocities of 25-200 nm/s (Gaska et al., 2020). Our *in vivo* data indicate that the resistive or brake forces produced by the microtubule-crosslinking activity of PRC1 are too low to oppose forces that drive spindle elongation (Fig 6). The speed of chromosome segregation and the distance between the segregated chromosomes mass in the presence of PRC1-3A are increased, compared with cells expressing PRC1 (Fig 6). This has also been reported for cells lacking PRC1 (Pamula et al., 2019; Vukusic et al., 2021), in which two half spindles became disconnected and were pulled apart. Because controlled microtubule sliding does not occur when PRC1 is not bound to the kinesin motors, the outwards spindle and cortical-generated forces are likely to dominate the system. We propose that PRC1 acts both as a break and signalling adaptor. Kinesin motors associate with PRC1 via the conserved ΦΦ motifs and either generate breaking forces on the spindle or recruit other signalling molecules to regulate cytokinesis. Kif4a, CENP-E, MKLP1 and MKLP2 all interact with PRC1 across species. Future work will address how these PRC1-interacting motors work collectively to completing cell division.

Failed cytokinesis is a hallmark of cancer cells, leading to chromosome instability. Fast growing polyploid cancer cells are particularly vulnerable to cytokinesis failure (McKenzie and D’Avino, 2016). Our work may open up opportunities to interfere with cytokinesis completion and induce cytokinesis failure in cancer cells as a therapeutic cancer target to increase chromosome instability and cell death (Lens and Medema, 2019).

## Methods

### Cloning

To assay the localization in cell culture of CENP-E subdomains, various constructs were generated from CENP-E transcript variant 1 (NM_001813.2) and cloned into pBABE-puro containing an N-terminal GFP tag and using restriction enzymes (Cheeseman and Desai, 2005). MBP-CENP-E was cloned into pMal-C2X (NEB). Bacterially-expressed constructs of GST-CENP-E were cloned in pET-3aTr (Tan, 2001). Mutagenesis was performed according to Quickchange mutagenesis protocols (Agilent). Mutants for CENP-E_2605-2701_ were synthesized using G-Blocks (IDT).

### Protein expression, purification and assays

All constructs for bacterial expression were transformed in *E. coli* BL21-CodonPlus (DE3)-RIL. Cultures were induced with 0.5 mM IPTG when OD_600_=0.6 for 4 hours at 25°C or overnight at 18°C for 18-20 hours. Cells expressing his_6_-proteins were re-suspended in lysis buffer (50 mM HEPES pH 7.5, 500 mM NaCl, 40 mM imidazole, 1 mM EDTA, 5 mM β-Mercaptoethanol) supplemented with 1 mM PMSF and cOmplete EDTA-free protease inhibitor cocktail (Roche) and lysed by sonication. The lysate was cleared by centrifugation (50 minutes, 22,000 RPM) in a JA 25.50 rotor (Beckman Coulter), filtered and loaded onto a HisTrap HP column (GE Healthcare). His_6_-tagged proteins were eluted in elution buffer (lysis buffer with 500 mM imidazole). Constructs containing a 3C protease cleavage site were incubated overnight in dialysis buffer (25 mM HEPES pH 7.5, 300 mM NaCl, 10 mM imidazole, 1 mM EDTA, 5 mM β-Mercaptoethanol) with 3C protease and then loaded onto a HisTrap HP column (GE Healthcare). MBP-CENP-E_2605-2701_ was purified using the same lysis buffer without imidazole and an MBP-Trap HP coloumn (GE Healthcare). For ITC, the MPB tag was cleaved overnight using Factor Xa (NEB) in dialysis buffer and loaded again on an MBP-Trap HP column. GST-proteins were purified as previously described (Legal et al., 2020). Recombinant proteins were then concentrated and loaded on a Superdex 200 Increase 10/300 GL (GE Healthcare) pre-equilibrated in size-exclusion chromatography buffer: 20 mM HEPES pH 7.5, 300 mM NaCl, 1 mM EDTA and 1mM DTT. Labelling of 647His_6_-GST-CENP-E was performed according to the manufacturer’s instructions with the AlexaFluor647 (A20173A, Invitrogen). Degree of labelling estimated as 1:3.

Full length CENP-E was expressed and purified as previously published (Craske et al., 2022). GFP-PRC1, PRC1 and PRC1 truncated fragments were purified as previously described (Subramanian et al., 2010). Porcine brain tubulin was purified as described (Castoldi and Popov, 2003) and stored in liquid nitrogen long term.

### Isothermal titration calorimetry

ITC experiments were carried out to determine the affinity and stoichiometry of PRC1:CENP-E constructs and Kif4A, known to bind PRC1. CENP-E peptides were synthesized by Lifetein, LLC. PRC1_1-168_, CENP-E constructs and GST-Kif4A_1133-1165_ were extensively dialysed into ITC buffer (20 mM HEPES pH7.5, 150 mM NaCl, 0.005% Tween-20, 0.5 mM TCEP); prior to the experiment to minimize heats of dilution upon titration. Peptides were directly diluted into ITC buffer. Protein concentrations were determined by absorption at 280 nm; extinction coefficients ε were as follows; PRC1_1-168_: 8480 M^-1^ cm^-1^, CENP-E_2605-2701_:6990 M^-1^ cm^-1^, GST-CENP-E_2605-2701_: 49850 M^-1^ cm^-1^. Peptide concentrations were determined by absorption at 214 nm; extinction coefficients ε were 22904 M^-1^ cm^-1^ for peptide 1 and 22983 M^-1^ cm^-1^ for peptide 2. For protein-protein ITC experiments, 1140, 224 and 224 μM PRC1_1-168_ were titrated into 56.1, 18.6 and 20.7 μM CENP-E_2605-2701_, GST-CENP-E_2605-2701_ and GST-CENP-E_2605-2701 2SD_ respectively at 25 °C in 16 aliquots: 1 of 0.5 μl followed by 15 x 2.5 μl. The concentration was calculated for the monomeric CENP-E constructs. For protein – peptide ITC experiments, 557 μM of PRC1_1-168_ (calculated for monomeric PRC1) was titrated into 15 μM peptide 1 or peptide 2 at 25 °C in 16 aliquots: 1 of 0.5 μl followed by 15 x 2.5 μl. The reference power was set to 3 μcal/s and syringe rotation 750 rpm. The enthalpy of binding was analysed with correction for heat of dilution using the software package provided by the instrument manufacturer (Auto-iTC200 microcalorimeter; Malvern Instruments). Data were fit to a simple binding model with one set of sites.

### Cell culture, immunofluorescence and microscopy

Stable clonal cells lines expressing GFP-PRC1 WT and GFP-PRC1 mutant (siRNA resistant and Cas9 resistant) were generated as described previously in cells expressing a guide RNAi targeting PRC1 and inducible expression of Cas9 (Cheeseman and Desai, 2005; McKinley and Cheeseman, 2017). To deplete PRC1, we treated the cells for 48 hours with an siRNA which target the 3’UTR of PRC1 (A-019491-15-0020, Horizon Dicovery). HeLa cells (93021013, Sigma Aldrich) were used and maintained in DMEM (Gibco) supplemented with 10% Tet-free FBS ( A4736401, ThermoFisher), 5% Penicillin/Streptomycin (Gibco) and 2.5 mM L-Glutamine at 37°C in a humidified atmosphere with 5% CO_2_. Cells are monthly checked for mycoplasma contamination (MycoAlert detection kit, Lonza). Transient transfections were conducted using Effectene reagent (Qiagen) or lipofectamine 3000 (Invitrogen) according to the manufacturer’s guidelines. Cells were washed in PBS and fixed in ice cold methanol or alternatively 3.8% formaldehyde in PHEM buffer (60 mM Pipes, 25 mM HEPES, 10 mM EGTA, 1 mM MgSO_4_, pH 7.0) for 10 minutes. For immunofluorescence, cells were incubated 5 minutes in pre-extraction buffer containing 22.6 nM _647_GST-CENP-E_2639-2671_ and fixed with 10 minutes cold methanol followed by 1 minute acetone treatment. Immunofluorescence in human cells was conducted as previously described using antibodies against tubulin (1:1000 anti-beta tubulin, mouse, T7816, Sigma OR 1:2000/1:1000 anti-alpha tubulin, rabbit, ab18251, Abcam) and PRC1 (sc-376983, Santa Cruz, 1:200 mouse) (McHugh et al., 2018). anti-CENP-E (Abcam, Ab5093; 1:1000 or 1:200), guinea pig anti-CENP-C (pAb; MBL PD030; 1:2000), secondary anti-rabbit Cy3 (1:400, Invitrogen) and anti-mouse Cy2 (1:800; Invitrogen). Hoechst 33342 (Thermo Fisher Scientific; H3570) was used to stain DNA. Images were obtained from a widefield Eclipse Ti2 (Nikon) microscope equipped with a Prime 95B Scientific CMOS camera (Photometrics), using a 100x objective (CFI Plan Apochromat Lambda, 1.49 N.A). 10-20 z-sections were acquired at 0.2-0.5 µm and presented as maximum intensity projections. For live-cell imaging, cells were transferred into a 35mm glass bottom viewing chamber (MatTek). Prior to imaging, cells were incubated for 5 minutes with CellMask orange (1:40000, Thermo Fisher Scientific) and 5 minutes with SPY650 (1:1000, Spirochrome). Cells were washed multiple times in L15 Leibowitz media (Gibco) supplemented with 10% FBS and 2.5 mM L-glutamine, prior to imaging on the widefield Eclipse Ti2 (Nikon) microscope equipped with a Prime 95B Scientific CMOS camera (Photometrics), using a 60x oil objective (CFI Plan Apochromat Lambda, Nikon, 1.3 N.A) and a heated chamber with CO_2_. Data were acquired for the 3 channels at 1minute interval with an optical spacing of 1.25 μm.

### Sample preparation for TIRF microscopy and TIRF Microscopy Imaging

For PRC1 and CENP-E, microtubule binding assays, 0.2 mg/mL GMPCPP (Jena Biosciences) microtubule seeds containing 7% rhodamine-tubulin (Cytoskeleton Inc., TL590M-B) were polymerized in BRB80 (80 mM PIPES pH 6.9, 1 mM EGTA, 1 mM MgCl_2_) for 1 hour at 37 °C, followed by centrifugation at 13,300 rpm for 10 minutes and then resuspended in BRB80. Anti–tubulin antibodies (Sigma, T7816) at a 1:10 dilution in BRB80 were first introduced to the chamber. Next, 40 μl of 1 % Pluronic F-127 (Sigma Aldrich) in BRB80 was washed through the chamber and incubated for 5 minutes. Chambers were then washed with 40 µL of BRB80, then 40 µL 1 mg/mL casein (Sigma Aldrich) before adding the final mixture of GMP-CPP microtubules and PRC1 and/or CENP-E in final assay mix at indicated concentrations (80 mM PIPES pH 6.9, 5 mM MgCl_2_, 1 mM DTT and an oxygen scavenger mix: 0.2 mg/ml glucose oxidase, 0.035 mg/ml catalase, 4.5 mg/ml glucose, and 0.1 % β-mercaptoethanol).

Sliding assays were carried out in flow chambers consisting of functionalised glass coverslips coated with PEG-biotin. Firstly, chambers were washed with BRB80. Next 40 μl of 50 μg/mL Neutravidin was washed through the chamber and left to incubate for 5 minutes. GMPCPP polymerised biotinylated tubulin (HiLyte647 labelled) was washed into the chamber and left for 5 minutes. Next, 50 μl of purified full-length PRC1 at 2.5 nM was added to coat the microtubules. This was left for 10 minutes. Chambers were then washed with BRB80, followed by flowing through with a final assay mix containing GMPCPP non-biotinylated microtubules, 2 mM ATP, 0.5 mg/ml casein, oxygen scavenger and CENP-E motor at indicated concentration (or buffer as a control). Microscopy was carried out immediately following this step. For microtubule sliding assays, images of free rhodamine-microtubules and immobilised HiLyte 647 biotinylated microtubules using the red and far-red channels respectively were taken every 2 seconds for a total of 20 minutes. Imaging was performed on a Zeiss Axio Observer Z1 TIRF microscope using a Zeiss 100 × NA 1.46 objective and either a Photometrics Evolve Delta electron-multiplying charge-coupled device camera or a Photometrics Prime 95B sCMOS camera controlled by Zeiss Zen Blue software.

### Image analysis

Quantification was done in Omero (OME, or ImageJ (National Institutes of Health)(Allan et al., 2012; Schneider et al., 2012). Linescans for measurement of intensity across the central spindle were generated for cells at a stage of cell division defined by taking the width of the spindle of around 10-12 pixels, visualized by tubulin staining with SiR dye (1:40.000, Spirochrome). For the measurement of the chromosome separation in telophase, the maximum distance between chromosome masses along the spindle axis was measured after maximum intensity projection of images. Cell stages were assessed by DNA morphology; telophase was distinguished from cytokinesis by the state of chromosome condensation and the shape of the cell; analysis was done in late anaphase and telophase cells, excluding cytokinesis. For quantification of midbody integrity, cells in cytokinesis were identified morphologically with a bundle of microtubules between neighbouring cells, marked by GFP-PRC1. For cells expressing PRC1 3A mutant in the absence of endogenous CENP-E, the bundle was generally partially or fully missing, or distorted and GFP-PRC1 was absent or on the remnant midbody, but chromosomes and microtubules could be seen between the two cells. Midbody was defined as a distinct PRC1 signal between the future daughter cells.

### Statistics and reproducibility

Statistical analyses were performed using GraphPad Prism 9.0. No statistical method was used to predetermine sample size.

### AlphaFold analysis

The PRC1:CENP-E dimeric complex structure was predicted using AlphaFold 2 in the multimer version.

### Data availability

All data and reagents supporting the findings of this study are available from the corresponding author on request. AU ideal to offer all data underlying the figures as Supplementary data.

## Author contributions

JW designed the project. AG, TL, BC and TM performed experiments. TL, BC, AGK performed data analysis and interpretation. JW wrote the manuscript. All authors approved the final draft of the manuscript.

## Acknowledgements

GFP-PRC1 and His-PRC1 were kind gifts from Tarun Kapoor (Rockfeller University). We are grateful to Iain Cheeseman (Whitehead institute for Biomedical Research, Cambridge, Massachussets, USA) for sharing the human HeLa inducible CRISPR/Cas9/PRC1-G2.2 knock-out (KO) cell line. We thank Owen Davies for help with AlphaFold2 and Liz Blackburn for help with ITC. We thank the Welburn lab for critical reading of the manuscript. J. W. is supported by a Wellcome Senior Research Fellowship (207430). This project was also supported by a COVID-reboot grant from the Royal Society of Edinburgh. The Wellcome Centre for Cell Biology is supported by core funding from the Wellcome Trust (203149), a Multi-User Equipment grant [101527] for the Edinburgh Protein Production Facility.

## Supplementary figures

**Supplementary figure 1:**
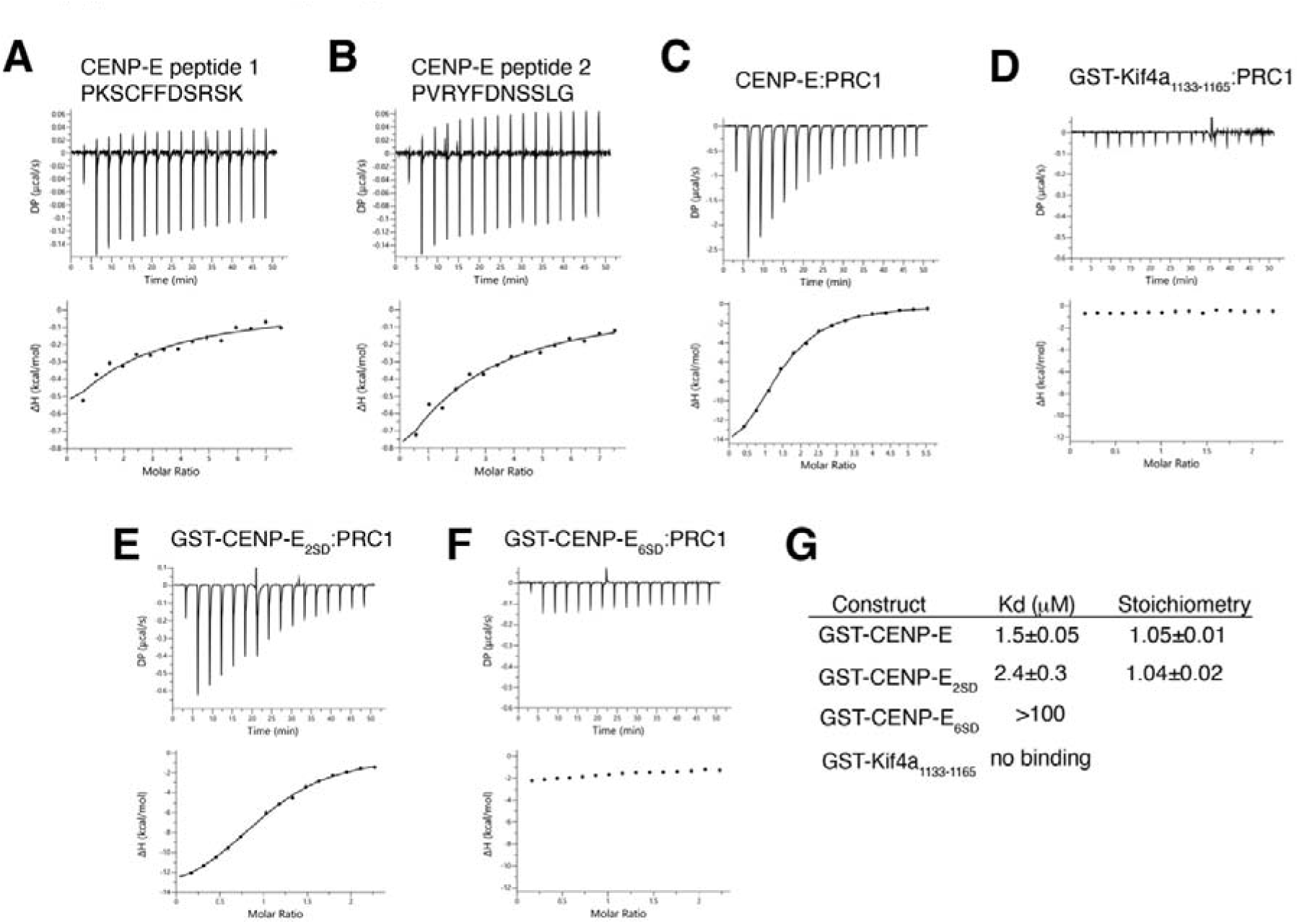
Characterization of the PRC1/CENP-E interaction. Characterization by isothermal titration calorimetry of binding between PRC1_1-168_ and CENP-E_2605-2701,_ GST-CENP-E_2605-2701,_ GST-CENP-E_2605-2701_2SD, GST-CENP-E_2605-2701_6SD and GST-Kif4A_1133-1165_. The y-axis indicates kcal/mole of injectant.

**Supplementary figure 2:**
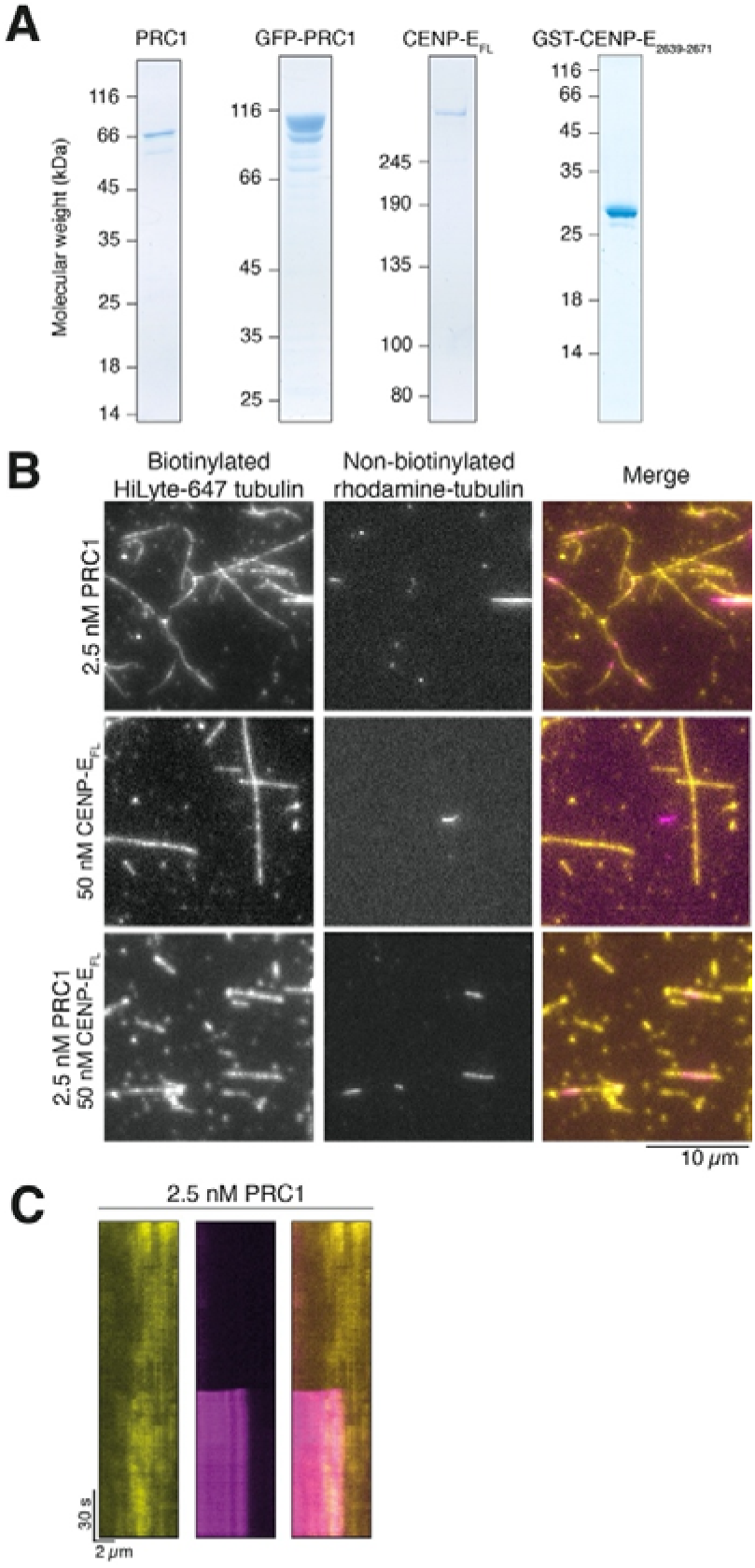
Human CENP-E_FL_ does not bundle microtubules in the presence of ATP. (A) Coomassie-stained gel showing purified His-PRC1, His-GFP-PFR1, Full-length CENP-E and GST-CENP-E_2639-2671_. (B) Bundling of microtubules by 2.5 nM PRC1 *in vitro*. Non-biotinylated rhodamine tubulin (Pink) and biotinylated HiLyte-647 tubulin (Yellow), scalebar 5 μm. (B) Crosslinking of microtubules by 2.5 nM PRC1, 50 nM CENP-E_FL_ or both *in vitro*. In the absence of PRC1 rhodamine-labelled microtubules in solution (pink) are not crosslinked to biotinylated surface bound Hilyte-647-labelled microtubules (yellow). Scalebar: 9 μm. (C) Kymograph showing cross-linking of a free microtubule to an immobilized microtubule after approximately 15 s of imaging.

**Supplementary Figure 3:**
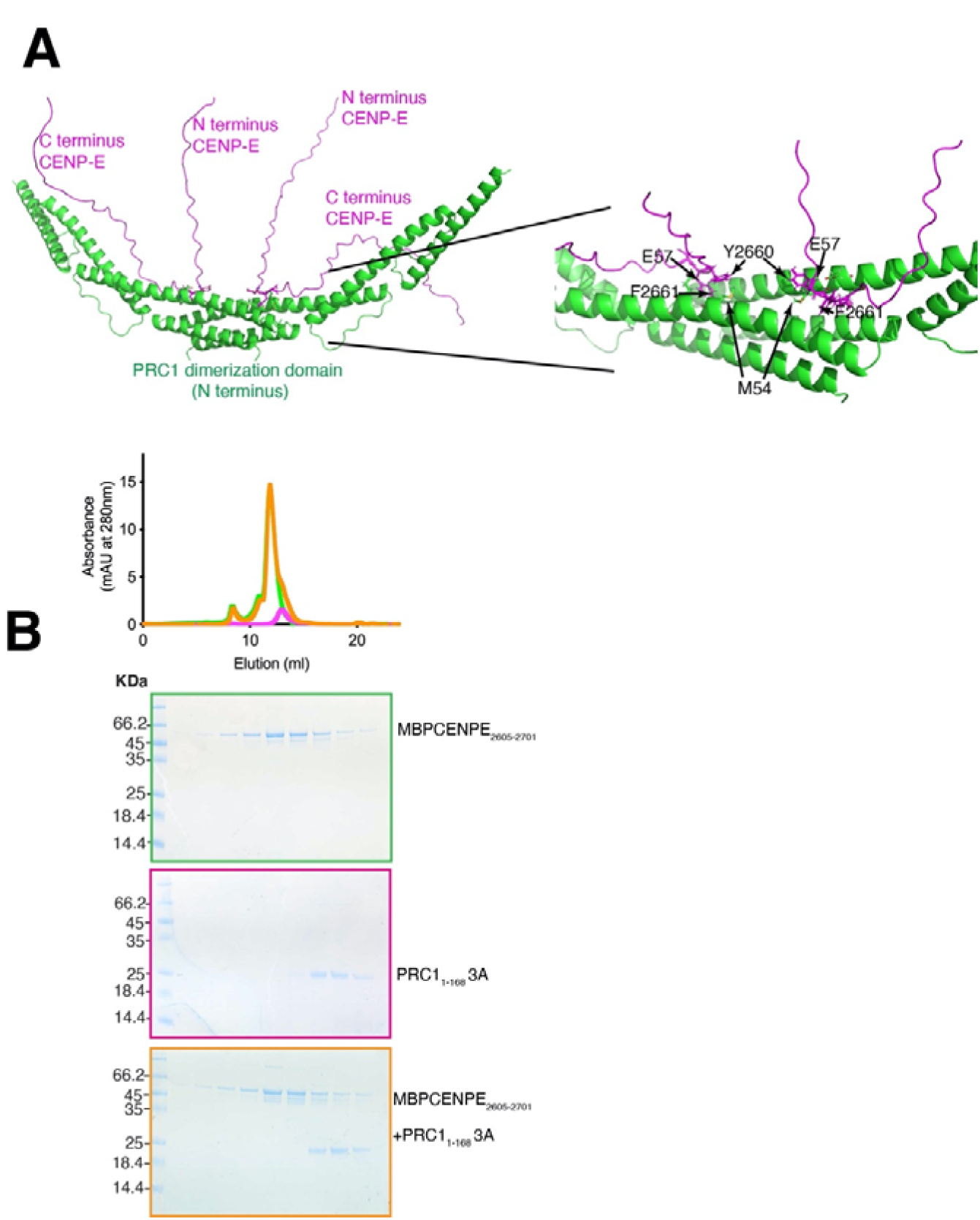
Molecular basis and regulation of the PRC1-CENP-E interaction. (A) AlphaFold2 prediction of the CENP-E/PRC1 interaction identifies the PRC1 residues important in CENP-E binding. (D) Size-exclusion chromatography elution profile of MBP-CENP-E_2605-2701_ (green), PRC1_1-168_ 3A (pink) and MBP-CENP-E_2605-2701_/PRC1_1-168_ 3A (orange). Bottom, coomassie-stained gel showing the size-exclusion chromatography profile of PRC1_1-168_ 3A (pink), MBP-CENP-E_2605-2701_ (green) and MBP-CENP-E_2605-2701_ /PRC1_1-168_ 3A (orange).

**Movie 1: Microtubule sliding in the presence of full-length CENP-E and PRC1.**

Free microtubules (magenta) slide past surface-immobilized microtubules (yellow) in the presence of 2.5 nM PRC1 and 50 nM full-length CENP-E.

